# Lack of evidence for sex differences in higher cognitive function in macaques

**DOI:** 10.1101/153593

**Authors:** Jamie R.J. Nagy, Christienne G. Damatac, Mark G. Baxter, Peter H. Rudebeck, Paula L. Croxson

## Abstract

Here we assessed whether higher cognitive function differed between male and female rhesus monkeys using tests of episodic memory and strategy implementation. We did not find any difference between males and females on behavioral performance or on analyses of grey matter volume of key regions. Our findings suggest that, at least where higher cognitive function in healthy monkeys is concerned, the sexes may not differ.

## Introduction

The incidence and symptom profiles of some psychiatric disorders differ markedly between men and women (Klein and Corwin, 2002). Sex differences in specific brain regions have been proposed to underlie these differences (McCarthy, 2015). Animal models may help bring clarity to biological mechanisms that mediate sex differences in brain structure and function. Critically, they may also provide insight into aspects of brain structure and function in which sex does not play a role. Unfortunately, the paucity of data on sex differences in behavior and brain structure in animal models makes it difficult to address the potential mechanisms by which sex influences psychiatric disorders. This problem is magnified because most preclinical animal studies use only males, limiting their generality (Clayton and Collins, 2014). This issue has resulted in recent NIH policies that sex be considered as a biological variable in all preclinical research, typically by including both males and females in all studies. However, the current pressure to balance sexes when carrying out animal studies may not always be practical, for example in non-human primate studies where numbers are often limited (Fields, 2014). Experimental designs that include both sexes without adequate statistical power to detect sex differences may also, inadvertently, contribute to problems with experimental reproducibility (Maney, 2016).Rhesus monkeys are uniquely suited as model organisms to address research questions related to higher cognitive functions in humans, such as those impaired in psychiatric illness, because their PFC is more similar to humans than any other animal available for research (Wise, 2008). The presence of sex differences in cognitive function and/or brain structure in this species could suggest that cognitive neuroscience studies in monkeys should include both sexes. Conversely, the absence would provide evidence that there are cognitive and brain functions that are not influenced by sex differences, meaning that studies of these specific kinds of functions, particularly in healthy animals, need not necessarily include both males and females. This is important as it would influence the factors to be considered in statistical power calculations and assessments of the number of animals to be used in future studies. Further, with the advent of advent of transgenic non-human primate models (Jennings et al., 2016) and the growing importance of non-human primate models in bridging the gap between genetic studies in animals and human neuropsychiatric disease, it is of critical importance to address which questions are best answered by non-human primate work.

Here, we addressed the question of sex differences on higher cognitive function by capitalizing on a large amount of behavioral data amassed over the past 10 years. Rhesus monkeys were tested under identical conditions on two tests of higher cognitive function. One task assessed episodic memory (object-in-place scene learning) and the other strategy implementation. Both depend on the integrity of specific subregions of prefrontal cortex (PFC) (Baxter et al., 2007; 2008c; 2009). We combined data across studies that were originally underpowered to consider sex differences, and re-analyzed them including sex as a factor.

In addition to studies considering the differences in behavioral effects between sexes, the question of whether there are structural differences in the brain between the sexes is equally prominent. However, studies of sex differences in structure of human brain regions have yielded divergent results (Ruigrok et al., 2014; Pintzka et al., 2015). To address this we analyzed high-resolution structural MRI data collected from a large cohort of monkeys to examine the effect of sex on the sizes of brain structures considered important for sex differences in the tasks studied here, and in human psychiatric disorders, namely the hippocampus, caudate, putamen and PFC.

## Materials and Methods

### Subjects

Data were acquired from rhesus macaque monkeys in the U.K. over a period of 10 years. All procedures were carried out in accordance with the U.K. Animals (Scientific Procedures) Act (1986). They were housed under a standard light/dark cycle (lights on 7:00AM - 7:00pm). All animals were housed in social groups of size ranging from 2 to 10, and group sizes varied in accordance with availability of other animals and ability to group successfully. Environmental enrichment was provided to all groups, and *ad libitum* water. Food was given once per day in accordance with behavioral testing schedules. Individual details of animals and housing are in the original publications.

### Behavioral tasks

#### Episodic memory behavioral task

We combined data from 56 rhesus macaque monkeys (45 male, 11 female) across eight studies (Baxter et al., 2007; Wilson et al., 2007; Baxter et al., 2008b; 2008c; 2009; Browning et al., 2010; Croxson et al., 2011; 2012) learning the object-in-place scene-learning task adapted from (Gaffan, 1994). Each unique scene had five elements: (1) the initial background color; (2) the location of ellipses and ellipse segments of random size, color, and orientation; and (3) the location, color, and symbol of a typographic character. Each scene further contained two small foreground objects placed in a constant location within each scene. For each scene, one foreground object was correct and rewarded and the second foreground object was incorrect and not rewarded.

#### Strategy implementation behavioral task

We combined data from 25 rhesus macaque monkeys (19 male, 6 female) across three studies (Baxter et al., 2007; 2009; Croxson et al., 2011) trained on the strategy implementation task described in (Baxter et al., 2007), a form of conditional discrimination learning. Monkeys were presented with pairs of clip art stimuli where for each pair, one stimulus represented a ‘persistent’ strategy (P), whereas the other stimulus represented a ‘sporadic’ strategy (S). To receive a reward using the P strategy, the monkey had to choose four consecutive persistent choice stimuli. Any time after this reward, a sporadic choice was rewarded on the next trial. However, sporadic choices were not subsequently rewarded until another reward from persistent choices had been earned. Therefore, in order to perform optimally in the strategy implementation task, monkeys had to learn to alternate between the P and S strategies upon receiving each reward (thus, an optimal trials/reward ratio was 2.5).

### MRI data acquisition and analysis

#### MRI data acquisition and preprocessing

We analyzed T1-weighted structural MRI images (acquired from the same 3 Tesla MRI scanner). Scans were acquired with a magnetization prepared rapid acquisition gradient echo (MPRAGE) sequence (no slice gap, 0.5 × 0.5 × 0.5 mm, temporal resolution = 2,500 ms, echo time = 4.01 ms, and 128 slices, flip angle = 7°) from male (n=12) and female (n=9) rhesus macaque monkeys (*Macaca mulatta*) aged between 3 and 5 years old at the time of scanning, under isoflurane anesthesia (1.0–1.8%). A full range of parameters was monitored and appropriate analgesia was given as necessary, as described previously (Mars et al., 2011). These data have been previously published (Mars et al., 2011; Sallet et al., 2011; O’Reilly et al., 2013). T1-weighted images were skull stripped using BrainSuite (Shattuck and Leahy, 2002) to skull-strip the original structural scans in order to remove non-brain material from further analysis and manually corrected each brain on FSL to verify accuracy. We additionally applied a bias field correction to the images using AFNI (Cox, 1996) to create a unifized image.

#### Hippocampal region of interest (ROI) tracing

Hippocampal ROIs were generated manually in FSLview (Smith et al., 2004) by an analyst blinded to the sex of each monkey, according to methods used by (Shamy et al., 2006). Structural scans were rotated along the long axis of the hippocampus to generate a straight region for tracing. Hippocampal ROIs were traced by hand on the rotated image starting in the coronal view and then checked in the sagittal and axial views. When tracing in the coronal view, we used the lateral ventricle as the dorsal boundary, with the ventral boundary being on the edge between gray and white matter below the subicular structures. After switching to the sagittal view, we extended the tracing rostrally to the lateral ventricle. At the caudal boundary, we included grey matter and excluded white matter. In subsequent sections, we avoided the amygdala rostrally by looking for differentiation in tissue contrast. We then realigned each rotated ROI of the left and right hippocampus onto the original structural space using the linear registration tool FLIRT (Jenkinson and Smith, 2001). Lastly, we generated a whole brain ROI of the skull-stripped structural scan in order to run a volumetric analysis on the whole brain and hippocampal regions. A second analyst blinded to the sex of each monkey then verified ROIs for accuracy.

#### Caudate, putamen, and PFC ROI tracing

Caudate and putamen ROIs and PFC subregion ROIs (rostral PFC (rPFC), ventrolateral PFC (vlPFC), dorsal PFC (dPFC), medial PFC (mPFC), and orbital PFC (oPFC) were generated from atlas regions (Paxinos et al., 2000; Markov et al., 2014) and nonlinearly registered to each monkey’s structural image using FNIRT in FSL. ROIs were then manually corrected by an analyst blinded to the sex of each monkey to exclude voxels in the ventricles and white matter and include grey matter voxels starting in the sagittal view, and then checked in the coronal and axial views. A second analyst blinded to the sex of each monkey then verified ROIs for accuracy.

### Statistical analyses

All statistical analyses were carried out in SPSS 22 (IBM, Armonk, NY). For the objectin-place-scene learning analysis, we performed a repeated measures ANOVA with a within-subject factor of trial and a between-subject factor of sex. For the strategy implementation data, we performed a one-way ANOVA with sex as a factor. For the anatomical analysis, we carried out multivariate analyses with repeated-measures ANOVA and subsequently *post* hoc analyses with independent samples t-tests assuming equal variances for (1) whole brain volume, and corrected volumes for (2) hippocampus, (3) caudate, (4) putamen, (5) rPFC, (6) vlPFC, (7) dPFC, (8) mPFC (9) oPFC. Details of statistics including effect sizes are reported in Supplementary Information. All graphs were created using GraphPad Prism 5 (GraphPad, San Diego, CA).

### Data availability

All relevant data are available on request from the authors.

### Results

Performance on the episodic memory task (object-in-place scene learning) was not influenced by sex (Figure 1a, b). Male (n=45) and female (n=11) monkeys did not differ in their performance (repeated-measures ANOVA, effect of sex, F_1,54_=0.089, p=0.767, Cohen’s d=0.14; sex by trial interaction, F_2.31,124.759_=0.366, p=0.724, d=0.14).

Similarly, there were no significant behavioral differences between male (n=19) and female (n=6) monkeys on the strategy implementation task for mean ratio of trials to rewards, a measure of efficient strategy use (Fig. 1c, one-way ANOVA, effect of sex, F_1, 23_=0.470, p=0.500, d=0.32). Taken together, these results indicate that in monkeys there are no differences in higher cognitive function, at least for the tasks used here.

**Figure 1.**
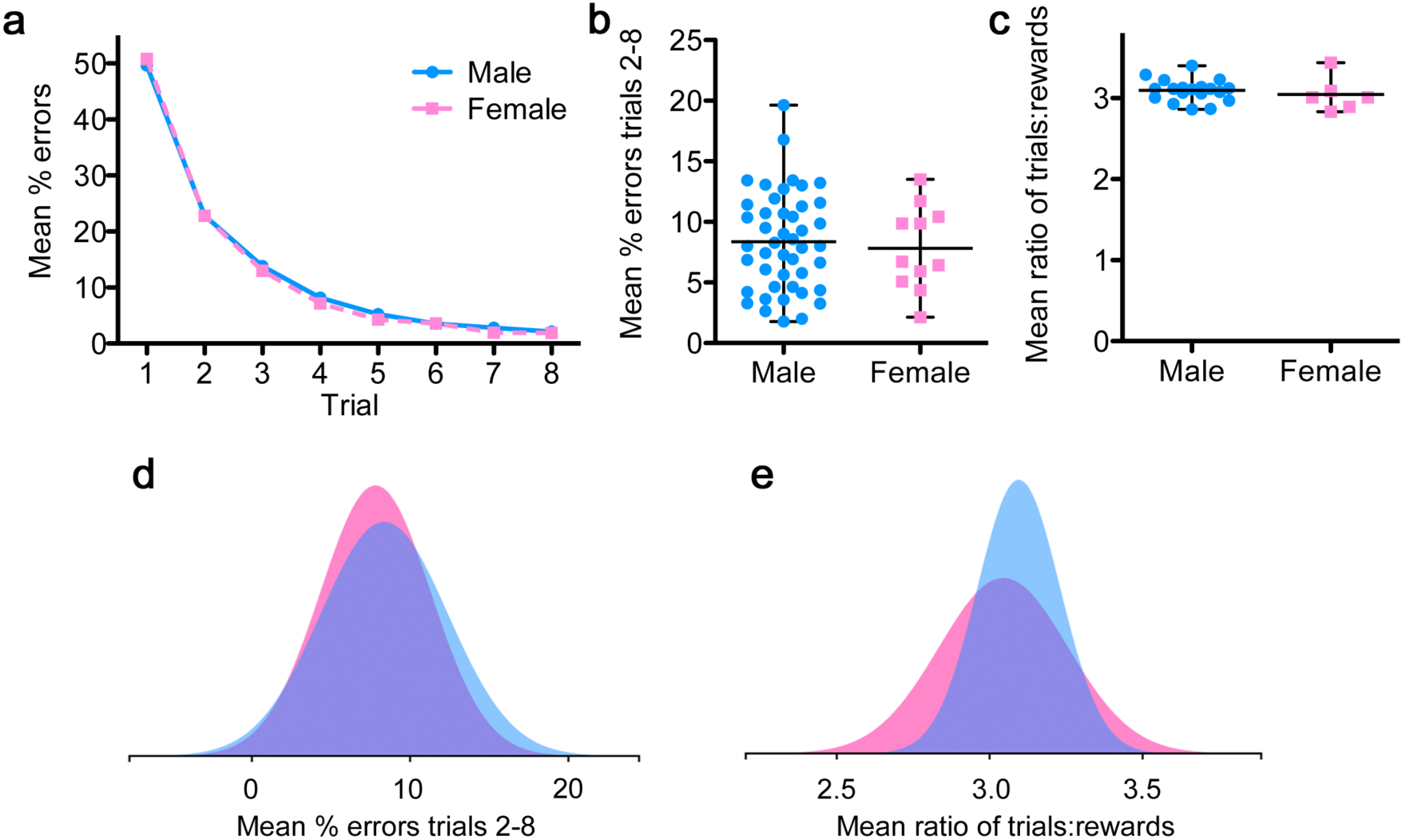
No behavioral differences between males and females. (a) Mean percent errors per trial depicting the learning curve for monkeys in the object-in-place scene learning task (n=45 males, n=11 females). (b) Mean percent errors for trials 2-8. A repeated-measures ANOVA revealed no main effect of sex (F1,54=0.089, p=0.767, d=0.14) and no interaction between sex and trial (F2.31, 124.759=0.366, p=0.724). Males: mean = 8.37, S.D. = 4.05; females: mean = 7.82, S.D. = 3.48. (c) Mean ratio of trials:rewards for the strategy implementation task (n=19 males, n=6 females). Optimal trials:rewards ratio is 2.5. A one-way ANOVA with sex as a factor revealed no main effect of sex (F1,23=0.470, p=0.500, d=0.32). Males: mean = 3.10, S.D. = 0.14; females: mean = 3.05, S.D. = 0.21. Black lines in (b) and (c) represent mean and range of data. Distributions for (d) object-in-place-scene learning and (e) strategy implementation showed 91% and 77% overlap respectively (generated with the sexdifferences.org tool from Maney et al. (2016)) (blue = male distribution, pink = female distribution, purple = overlap).

MRI volumetric analyses in *a priori* regions of interest (hippocampus, caudate, putamen, and PFC subregions; Figure 2a, c) were corrected for overall brain volume as this differed between the sexes (see Supplementary Information). When correction for whole-brain volume was used (regional volume as a proportion of whole-brain volume), there was no evidence of sex differences in regional volume (multivariate analysis with repeated-measures ANOVA: no main effect of sex (F_1,19=_0.244, p=0.627) and no interaction between sex and region (F_7,133_= 1.151, p=0.336) (Figure 2b, d). If, however, we lowered the threshold for this analysis one region repeatedly emerged, medial PFC (mPFC). This area was larger in females than in males and despite failing to reach statistical significance, showed the largest effect size (d=1.29).

**Figure 2.**
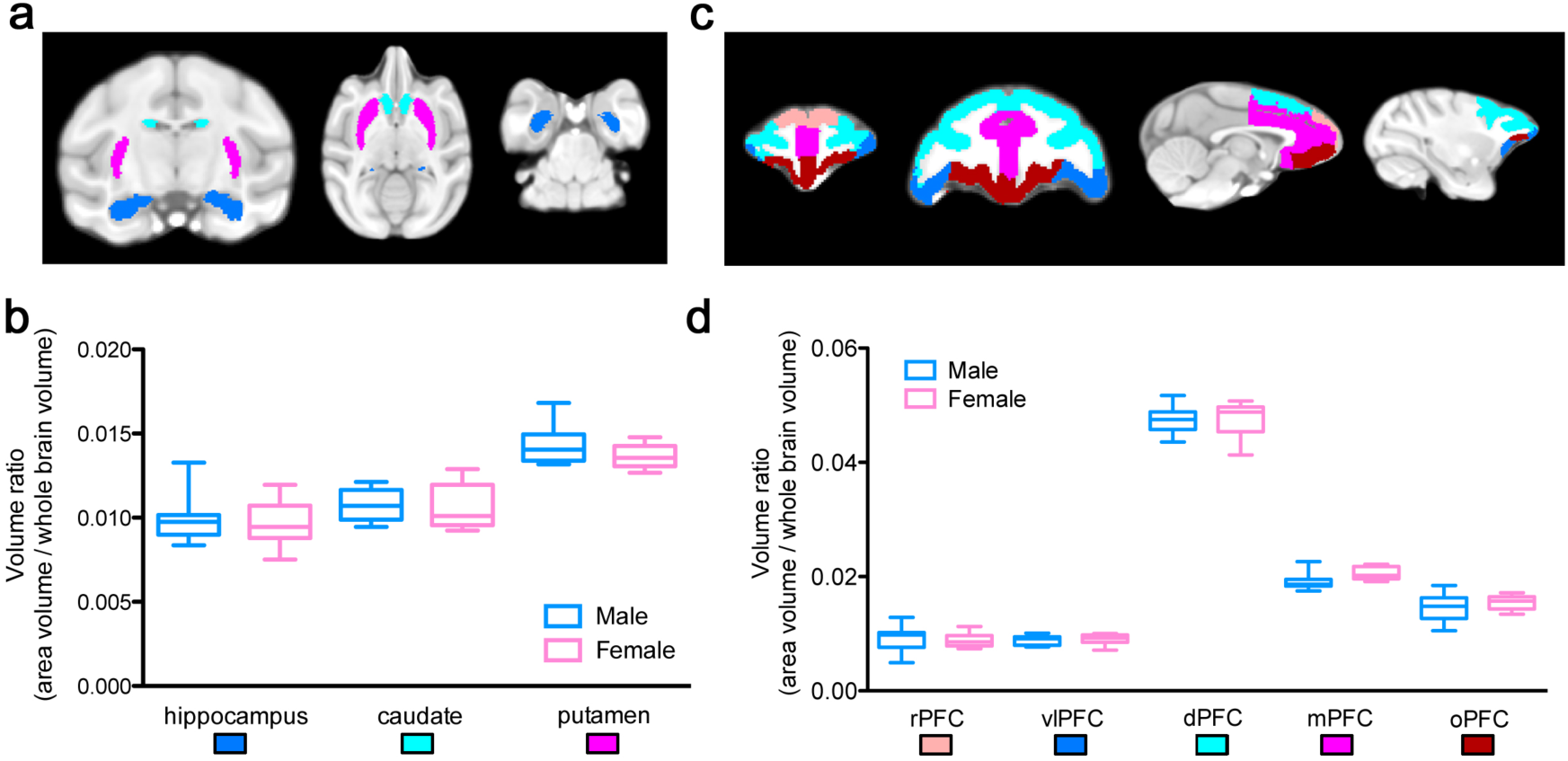
No differences in regional volume (corrected for whole brain volume) between males and females (n=12 males, n=9 females). (a) Subcortical regions of interest shown on MNI standard brain. (b) Volume ratio in the hippocampus (blue), caudate (cyan) and putamen (magenta). (c) Cortical regions of interest shown on MNI standard brain. (d) Volume ratio in the rostral prefrontal cortex (rPFC; pink), ventrolateral prefrontal cortex (vlPFC; blue), dorsal prefrontal cortex (dPFC; cyan), medial prefrontal cortex (mPFC; magenta) and orbital prefrontal cortex (oPFC; red). A repeated-mesures ANOVA with region as the within-subjects factor and sex as the between-subjects factor revealed no main effect of sex (F1, 19=0.244, p=0.627) and no interaction between sex and region (F7, 133=1.151, p=0.336). Centre line of box plots is mean, box is inter-quartile range, bars are data range.

There was a significant difference in whole brain volume in mm^3^ between the sexes (n=12 males, n=9 females), with males having larger brains overall (independent samples t-test, t(19)=2.734, *p*=0.013; d = 1.21) (**Figure 3a)**. *Post hoc* tests revealed no significant effect of sex in any region when Bonferroni correction was applied for multiple comparisons (i.e. yielding a significance threshold of *p*<0.006) (**Table 1)**. However, one region – the mPFC – did approach significance (*p*=0.009). As well as main statistical effects for each test, we report here effect sizes (*Cohen’s* d) for each test as a measure of whether we have sufficient statistical power to assess the group difference in each case, since it is not appropriate to carry out a full power analysis *post hoc*.

**Table 1.**
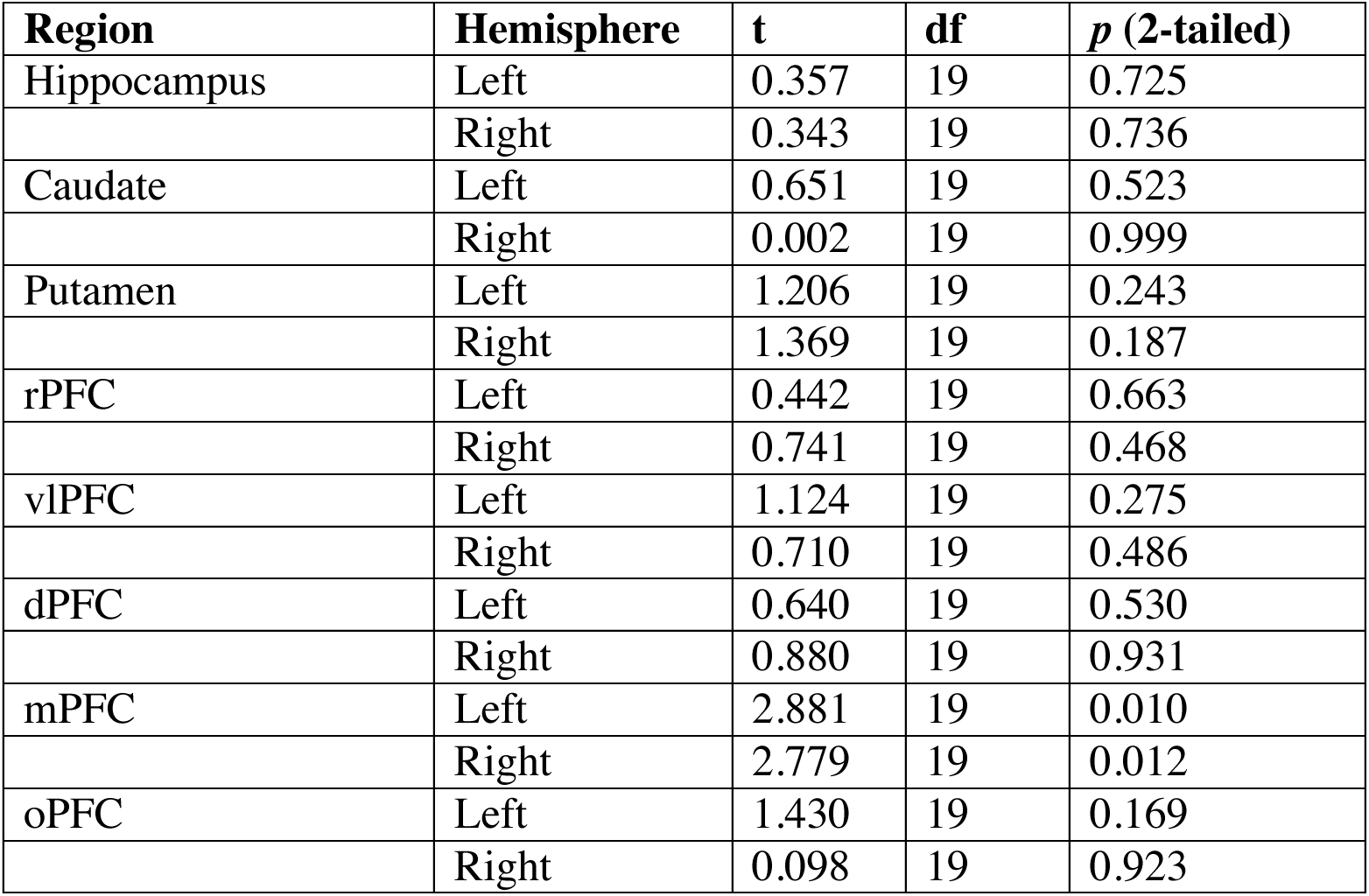
*Post hoc* test results for each subcortical and cortical region.

We also assessed hemispheric differences between the sexes (also corrected for whole-brain volume). A repeated-measures ANOVA with within-subjects factors of region and hemisphere, and between-subjects factor of sex revealed no significant interaction between sex, region and hemisphere (F_7,133=_1.151, p=0.335). Results of independent samples t-tests are shown in **Table 2** below and in **Figure 3b** and **c**. No results were statistically significant at *p*<0.003 (Bonferroni-corrected for multiple comparisons). However, the mPFC again had a very low *p* value.

**Figure 3.**
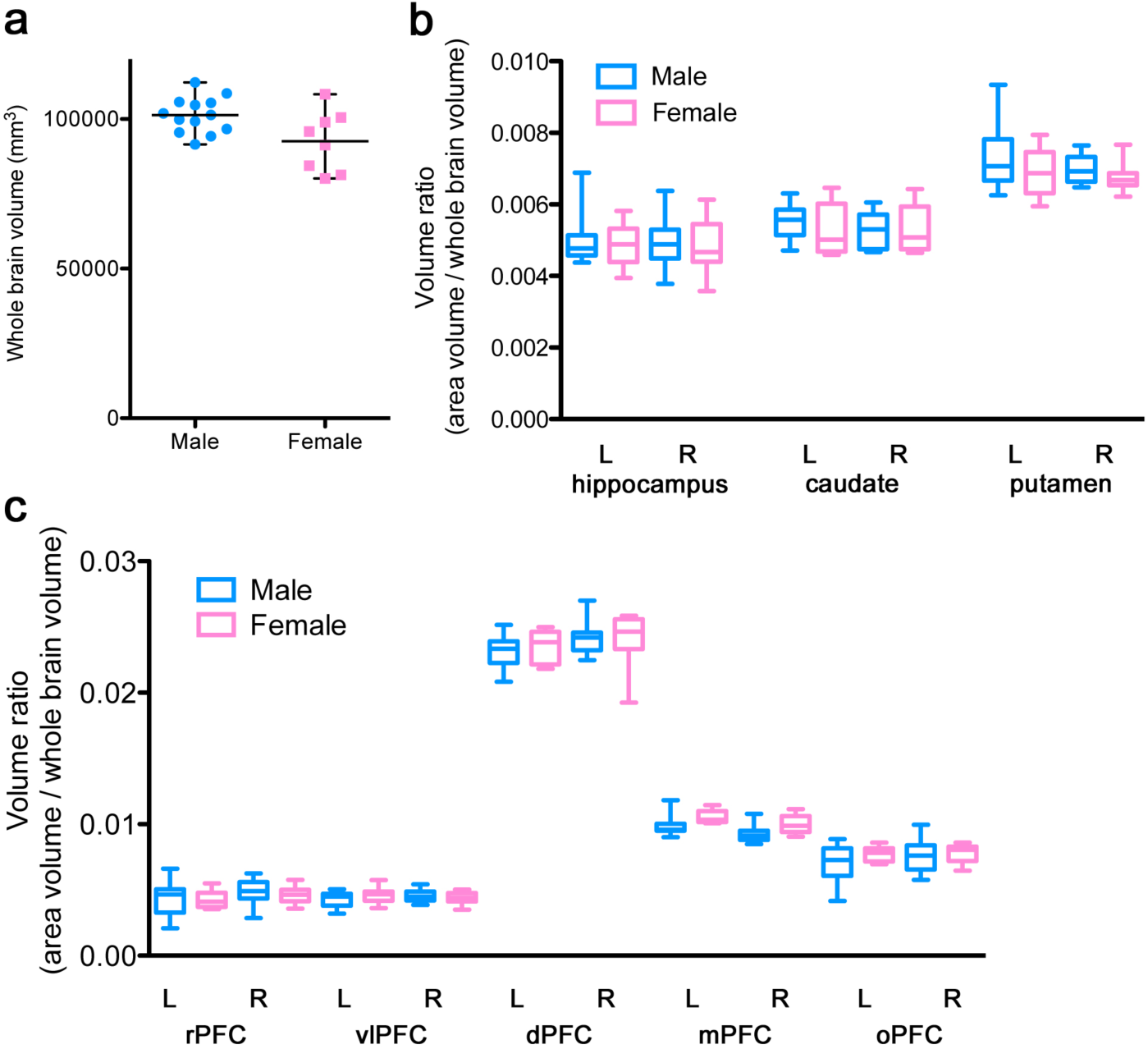
Whole brain and regional volumes assessed by hemisphere. (a) Male brains are significantly larger than female brains. Black lines represent mean and range of data. However, there were no differences in hemispheric regional volume between males and females when corrected for whole brain volume between males and females in (b) Subcortical regions of interest, (c) Prefrontal cortex subregions: rostral prefrontal cortex (rPFC), ventrolateral prefrontal cortex (vlPFC), dorsal prefrontal cortex (dPFC), medial prefrontal cortex (mPFC) and orbital prefrontal cortex (oPFC). Centre line of box plots is mean, box is inter-quartile range, bars are data range.

**Table 2.**
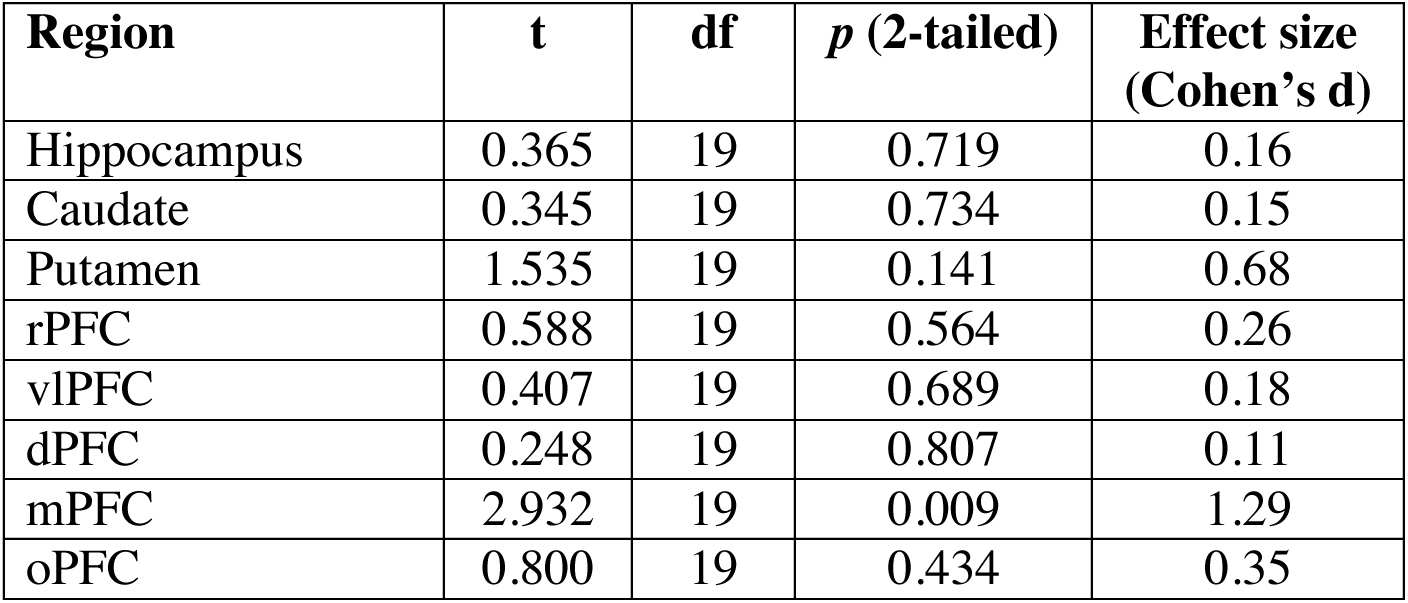
*Post hoc* test results for each subcortical and cortical region divided by hemisphere.

A whole-brain deformation-based morphometry analysis similarly found no differences between the sexes when corrected for multiple comparisons (either using threshold-free cluster enhancement or voxel-wise analysis, no voxels at *p*<0.05). However, using a stricter threshold but not correcting for multiple comparisons (voxel-wise p<0.005, cluster size > 5mm^3^) revealed bilateral clusters in the dorsal anterior cingulate cortex in the female > male contrast and bilateral superior temporal sulcus, rostral prefrontal cortex and cerebellum, as well as unilateral ventral premotor cortex (Figure 4). The dorsal anterior cingulate cortex may be involved in a sex difference during autobiographical memory processing in humans (Compère et al., 2016).

**Figure 4.**
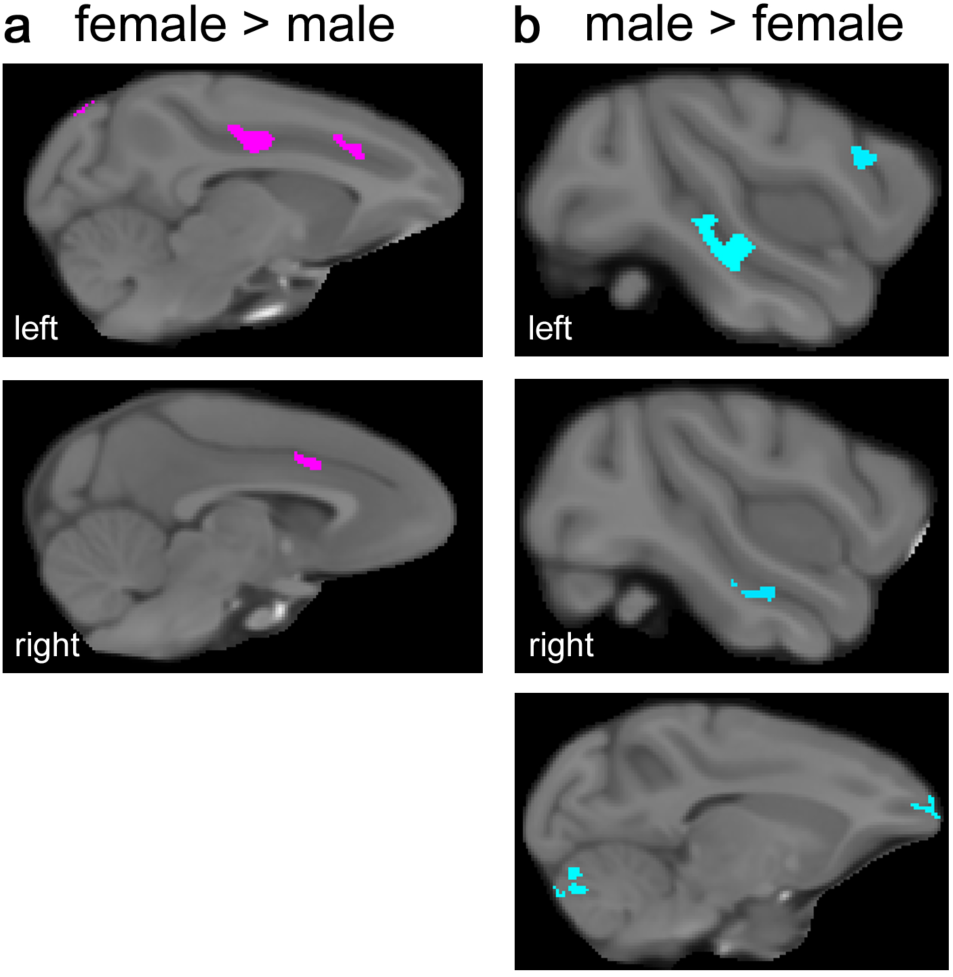
Whole-brain volumetric analysis. Despite no significant differences in volume when corrected for multiple comparisons, at a lower threshold (p<0.005, cluster size >5mm3), there were specific regional differences in brain volume between males and females. (a) Female > male: bilateral dorsal anterior cingulate cortex. (b) Male > female: bilateral superior temporal sulcus, rostral prefrontal cortex and cerebellum, as well as unilateral ventral premotor cortex.

## Discussion

In summary, we did not find sex differences in performance of PFC-dependent tasks of higher cognitive function in rhesus monkeys (Figure 1). This indicates that sex can be ruled out as a factor potentially influencing behavioral results obtained with these tasks, and potentially others assessing higher cognitive function in monkeys.

Males and females differ in the prevalence, age of onset, and symptomatology of many neuropsychiatric conditions (see (Ruigrok et al., 2014) for review). Dysfunction within brain regions involved in higher cognitive function, particularly the hippocampus and PFC, is apparent in neuropsychiatric disorders including schizophrenia, depression, and post-traumatic stress disorder (Godsil et al., 2013). The tasks we used here test episodic memory and strategy implementation, two cognitive tasks that are critically dependent on the function of PFC subregions (Baxter et al., 2007; 2008a; 2009) and the hippocampus (Browning et al., 2012). Both are cognitively demanding and rely on the integration of information across time, a function that is highly dependent on the PFC (Browning and Gaffan, 2008).

Complementary to our behavioral findings, there were no significant differences between the sexes in the volumes of the PFC, hippocampus, caudate or putamen (Figure 2). This is parsimonious with the findings of more recent human MRI analyses and meta-analyses that found no differences in the volume of brain regions including the hippocampus, caudate and putamen between males and females when taking whole-brain volume into account (Pintzka et al., 2015; Tan et al., 2015). The behavioral tests used here, which model the function of homologous human brain areas (Gaffan, 1994; Baxter et al., 2009), and the brain regions essential for them, particularly those in PFC, may represent a point in evolutionary divergence when specialized higher cognitive functions emerged that may operate outside of fundamental sex differences (Wise, 2008). We note that other measures such as synaptic patterning, firing rates, neurochemical identity, cell density, projection size, and transmitter release rates may provide alternative indices of brain function to the MRI data we present here, and future studies in this area may shed more light on the subject.

We do not consider that our findings are at odds with the extensive literature describing sex differences in brain development, hormonal receptor distribution, anatomy of sexually dimorphic areas, and behavioral differences in cognitive functions that are not reliant on higher-order association cortex (Cahill, 2006; de Vries and Södersten, 2009). For the assessment of behavioral performance and brain structure volume conducted here, fundamental differences in these measures are not present, or at least are so small that they are of no practical significance. This is entirely consistent with findings that the male and female brain are not categorically different, more on a spectrum (Joel et al., 2015).

Notably, it took many years to accrue enough data for the present study to be adequately powered to study sex differences. On the basis of these data we conclude that it is unnecessary to include adequately-sized and balanced groups of both sexes in studies of higher cognitive function in monkeys (Fields, 2014), particularly as unknown sex differences may compromise the generality of the results. Further, while including subjects of both sexes may not increase variance in performance as has been reported in rodents (McCarthy, 2015), and taking into consideration the absence of sex differences reported here, such a requirement would likely impact the welfare of the subjects (e.g. halving potential social group sizes). In non-human primate studies, which necessarily have small group sizes, it is important to consider whether pressure to include additional animals in order to include subjects of both sexes is justified and consistent with the reduction, refinement and replacement of animals used in research. Therefore the decision to include subjects of both sexes in studies in monkeys should be taken with care. Preclinical research has much more to do to address the underlying basis of sex differences that have already been identified in human psychiatric illness. Only by specifically designing studies to elucidate the mechanisms of these differences will we be able to uncover the fundamental biological importance of sex.

## Acknowledgements

We used data from the following publications, published over the last 10 years, all cited in the text, and including the following authors, to whom we are grateful: Philip G Browning, David Gaffan, Diana Kyriazis, Charles RE Wilson and Anna S Mitchell.

